# Poor availability of context-specific evidence hampers decision-making in conservation

**DOI:** 10.1101/2020.02.13.946954

**Authors:** Alec P. Christie, Tatsuya Amano, Philip A. Martin, Silviu O. Petrovan, Gorm E. Shackelford, Benno I. Simmons, Rebecca K. Smith, David R. Williams, Claire F. R. Wordley, William J. Sutherland

## Abstract

Evidence-based conservation relies on robust and relevant evidence. Practitioners often prefer locally relevant studies whose results are more likely to be transferable to the context of planned conservation interventions. To quantify the availability of relevant evidence for amphibian and bird conservation we reviewed Conservation Evidence, a database of quantitative tests of conservation interventions. Studies were geographically clustered and found at extremely low densities - fewer than one study was present within a 2,000 km radius of a given location. The availability of relevant evidence was extremely low when we restricted studies to those studying biomes or taxonomic orders containing high percentages of threatened species, compared to the most frequently studied biomes and taxonomic orders. Further constraining the evidence by study design showed that only 17-20% of amphibian and bird studies used robust designs. Our results highlight the paucity of evidence on the effectiveness of conservation interventions, and the disparity in evidence for local contexts that are frequently studied and those where conservation needs are greatest. Addressing the serious global shortfall in context-specific evidence requires a step change in the frequency of testing conservation interventions, greater use of robust study designs and standardized metrics, and methodological advances to analyze patchy evidence bases.

## Introduction

Tackling the biodiversity crisis with limited resources requires efficient and effective conservation action (Dirzo et al., 2014; Sutherland, Pullin, Dolman, & Knight, 2004). To inform which conservation actions (‘interventions’) are effective and which are not, we need a large and robust evidence base, ideally including large numbers of studies (replication of evidence; Fig.1A) with high internal validity (quality; Fig.1A) and external validity (relevance; Fig.1A). However, the limited resources available for conservation research mean that the evidence base for conservation is geographically and taxonomically biased (Christie, Amano, Martin, Petrovan, et al., 2019; Fazey, Fischer, & Lindenmayer, 2005; Hickisch et al., 2019; Spooner, Smith, & Sutherland, 2015). This is likely to limit the quality and relevance of evidence and impair effective decision-making (Cook, Possingham, & Fuller, 2013). Quantifying the availability of relevant, reliable studies is necessary to understand the strength of evidence upon which decisions are made, and to prioritize research on the effectiveness of conservation interventions.

**Figure 1.**
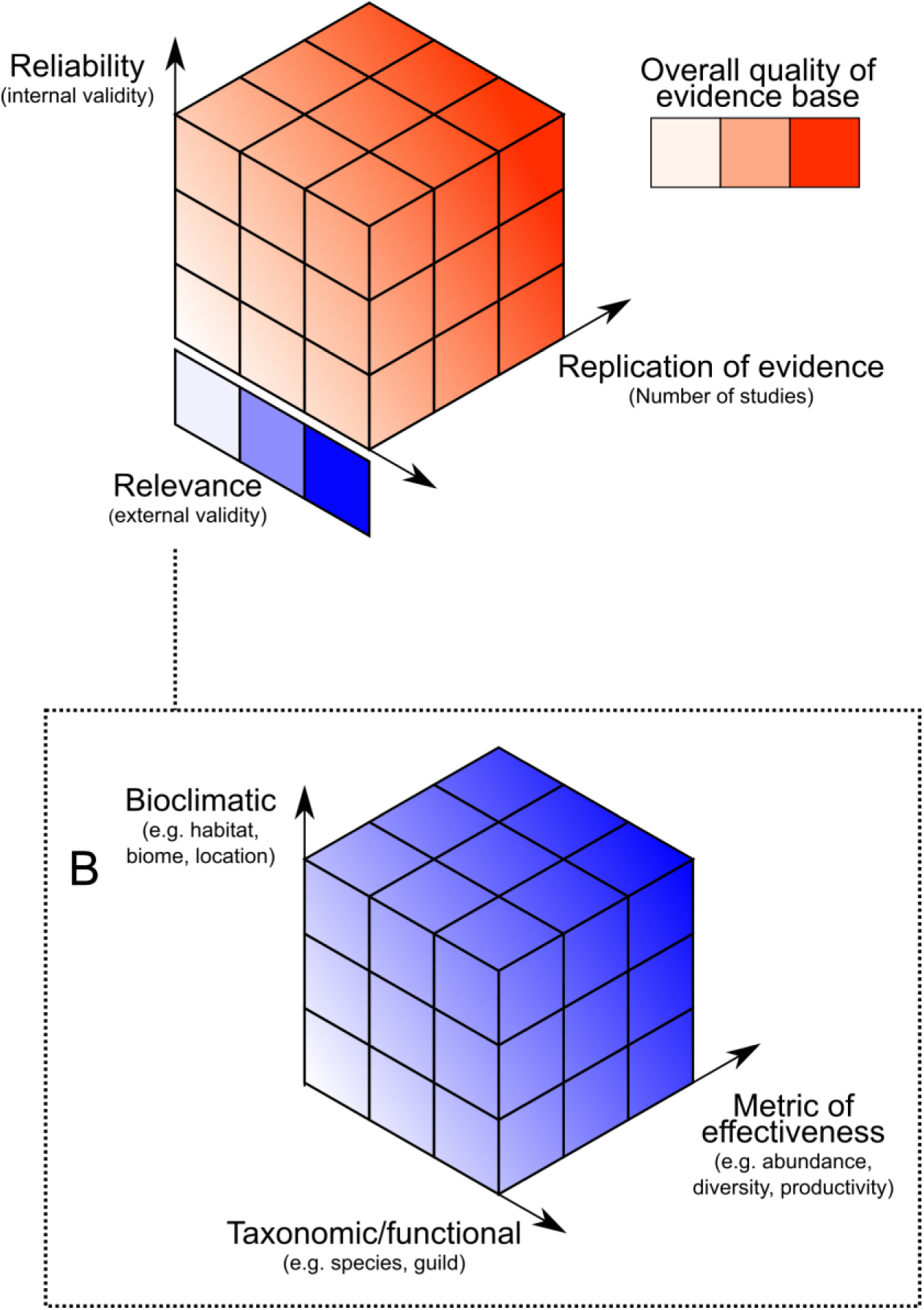
Framework of the desirable aspects of an ideal evidence base (stronger colors = more desirable). Fig.1A shows the three major desirable factors that an evidence base should have; large replication of evidence that is highly reliable (high internal validity) and highly relevant (high external validity). Fig.1B refers to the three dimensions that we will focus on that influence the overall relevance of evidence: i) bioclimatic (e.g., the study system), ii) taxonomic/functional (the study taxa) and iii) effectiveness measure (how you define and measure conservation success).

Practitioners and policymakers typically prefer to base their decisions on studies that are relevant (i.e., with high external validity; Fig.1) to their local context (Addison, Cook, & de Bie, 2016; Cook & Sgrò, 2017; Geijzendorffer et al., 2017). Using context-specific studies as evidence helps to ensure that results are likely to be repeated if the intervention is implemented again. The relevance of conservation studies to a given context will span multiple dimensions, including: (i) bioclimatic (i.e., similarity between habitats or regions); (ii) taxonomic/functional (i.e., similarity between taxa in terms of ecological function or taxonomic groups); and (iii) which metric was used to quantify the effectiveness of an intervention (i.e., the response variables or metrics of interest; Fig.1B). Other dimensions may also be important, such as the similarity between a study’s and a practitioner’s socioeconomic and political contexts, but we focus on the three dimensions above.

The first of these dimensions - bioclimatic relevance - refers to the similarity between the study ecosystem and the practitioner’s ecosystem (Fig.1B). The second dimension - taxonomic/functional relevance - concerns the similarity between the focal taxa of a study and the taxa of interest to the practitioner (Fig.1B). Together, these determine the ecological similarity between study and practitioner local contexts. This is vital because responses to interventions will vary between ecosystems and taxa. For example, the effectiveness of artificial nest boxes varies between different countries and habitats (Finch et al., 2019), while the effectiveness of translocation for New Zealand robins (*Petroica australis*) is unlikely to be relevant to a practitioner translocating Kakapo (*Strigops habroptila*). Practitioners who are interested in broader functional groups (e.g., seed dispersers or pollinators), taxa (e.g., birds, amphibians), or even whole ecosystems, may focus more on the functional relevance rather than taxonomic similarity of studied species.

The third dimension of relevance is the metric used to measure the effectiveness of an intervention. Practitioners may be interested in different responses to interventions depending on their focus (e.g., species or ecosystem-level responses) and effectiveness may vary depending on the metric used (Capmourteres & Anand, 2016; Marshall, Wintle, Southwell, & Kujala, 2019). For example, at the ecosystem-level, the effectiveness of bird boxes may be measured using the species richness or diversity of birds using them (Caine & Marion, 1991), while at the species-level, the number of individuals (Brawn & Balda, 1988), fledglings (Male, Jones, & Robertson, 2006; Purcell, Verner, & Lewis W, 1997), or brood size (Browne, 2006) may be measured. Similarly, the effectiveness of road mitigation interventions (e.g., tunnels or bridges) may be measured by the numbers of individuals of different species using the structures, but could also be measured in terms of levels of road mortality (Helldin & Petrovan, 2019). Therefore, the type of metric used by studies to measure effectiveness can have a major influence on the relevance of evidence.

The reliability of an evidence base - the internal validity of its studies - ultimately determines the overall quality of the evidence base and depends to a large extent on study design (Christie, Amano, Martin, Shackelford, et al., 2019; De Palma et al., 2018; Spake & Doncaster, 2017). As the conservation evidence base contains a wide variety of study designs (De Palma et al., 2018), there is likely to be variation in the reliability of inferences that can be drawn (Christie, Amano, Martin, Shackelford, et al., 2019). This variation may lead scientists to make misleading recommendations to practitioners, ultimately reducing the effectiveness of conservation practice, and making it difficult for decision-makers to weigh the strength of evidence provided by different studies.

The replication of evidence - the number of studies in the evidence base - is also important as greater numbers of studies demonstrating repeatable and reproducible effectiveness will give us greater confidence in the overall strength of the evidence. Decision-makers should rightly be wary of basing decisions on a low number of studies where reproducible effectiveness has not been or cannot be demonstrated - particularly given the current reproducibility crisis (Begley & Ioannidis, 2015; Nosek & Errington, 2017; Open Science Collaboration, 2015). However, the overall number of studies is not the only indicator of the strength of the evidence, since studies with low internal validity (e.g., poor study designs) and/or external validity (i.e., low relevance) may not constitute reliable evidence. Currently, we have a poor quantitative understanding of the availability of relevant and reliable studies in the conservation literature.

In this study, we assess whether studies testing conservation interventions are distributed across different contexts (bioclimatically, taxonomically, and by the metric used to measure effectiveness) in ways that reflect the needs of conservation. We also quantify other desirable aspects of the evidence base for conservation in terms of the quantity and quality of available studies; i.e., the number of studies that have tested different conservation actions, and how many of these use robust study designs.

## Methods

### Conservation Evidence database

We assessed the availability of relevant evidence for conservation practice using Conservation Evidence, a database of 5,525 publications as of January 2020 (Conservation Evidence, 2020a) that have quantitatively assessed the effectiveness of conservation interventions. Interventions are defined as management actions that a practitioner may undertake to benefit biodiversity (see Sutherland et al. (2019) for detailed methods). When we refer to the number of studies per intervention, we refer to the number of different tests of interventions - single publications may report multiple tests of different interventions. We assessed the availability of evidence for amphibians and birds based on synopses compiled in 2014 (n=419 studies; Smith & Sutherland, 2014) and 2012 (n=1,232 studies; Williams et al., 2013), respectively. More recent publications will obviously have increased the evidence base, but the broad patterns we quantify are unlikely to have changed in the intervening years. We excluded meta-analyses or systematic reviews from our analyses as these typically cannot be attributed to a particular local context (e.g., biome or taxon). We also only included interventions for which studies were present in the database. Since 32% (n=33) of interventions for amphibians and 25% (n=80) of interventions for birds had no associated studies in the database (i.e., were untested or tests were unpublished) or only included reviews or meta-analyses, the following analyses are likely to be an optimistic assessment of the availability of evidence in conservation. We used R statistical software version 3.5.1 (R Core Team, 2019) for all analyses.

### Local availability of studies by geographical distance

To calculate the average availability of studies within a certain distance of a given practitioner’s location, we generated 1,000 regularly spaced coordinates across certain parts of the world. For amphibians, we spaced these coordinates over the combined extent of all amphibian species ranges (IUCN, 2019) as this represents the possible range of locations in which a practitioner might conduct an intervention to conserve amphibians. For birds, we spaced these coordinates across the world’s terrestrial land masses (using “OpenStreetMap” 2019; see Appendix S1 for maps of coordinates) since although the combined distribution of all bird species is almost global, most practitioners are likely to conduct interventions to conserve birds terrestrially. Although non-terrestrial interventions are carried out by practitioners, the vast area covered by the ocean would severely underestimate the availability of studies to a practitioner’s likely location. 19 non-terrestrial interventions for birds were found in the database (e.g., ‘use streamer lines to reduce seabird bycatch on longlines’ or ‘use high-visibility mesh on gillnets to reduce seabird bycatch’) containing 33 studies in total - these were still included in our analysis as these studies tended to be conducted within close proximity to a terrestrial landmass (i.e., coastal).

We then calculated the Great Circle Distance from each study to each coordinate (see Appendix S1 for details), binning distances into a series of categories (100 km, 1,000 km and then every 1,000 km up to and including 19,000 km). We also calculated the ‘Global Mean’, which is the mean number of studies per intervention in the entire database - equivalent to approximately 20,000 km at the equator, the maximum distance separating any two coordinates. We then calculated the mean number of studies within each distance bin across all coordinates, as well as the number of studies that used different categories of study designs: i) any design, ii) Before-After (BA), Control-Impact (CI), Before-After Control-Impact (BACI) or Randomized Controlled Trial (RCT); iii) CI, BACI or RCT; iv) BACI or RCT designs (see Methods in Christie, Amano, Martin, Petrovan, et al. 2019 for definitions of each design).

We then repeated this analysis using the same number of coordinates (n=1,000), but this time by randomly selecting coordinates from amphibian and bird studies in the database (sampling with replacement from amphibian studies as there were fewer than 1,000). Using both approaches provided likely upper and lower bounds of evidence availability: regular coordinates likely underestimated the availability of evidence to practitioners, giving equal weighting to locations where conservation interventions are unlikely to occur (e.g., Antarctica) and those that are more intensively managed (e.g., Europe). In contrast, using locations from existing publications will likely overestimate study availability as this assumes that practitioners only conduct interventions in locations where they have previously been tested.

We compared the results of the first analysis (regularly spaced coordinates) to the expected patterns we would observe if studies were regularly distributed. We did this by generating equal numbers of regularly spaced coordinates (‘expected studies’) as the number of amphibian and bird studies (419 and 1,232 coordinates, respectively) using the same methods and shapefiles as before. We then calculated the mean number of these ‘expected studies’ within each distance bin.

### Context-specific availability of studies

To quantify the amount of relevant and robust evidence on the effectiveness of different conservation interventions, we required metadata that described each study’s local context and study design. By adapting previously described methods (Christie, Amano, Martin, Petrovan, et al. 2019; Appendix S2), we extracted the biome, taxonomic order and reported metric type used by each study (to quantify the number of relevant studies), as well as the broad category of study design used (to quantify the number of robustly designed studies). When metric metadata was extracted, we grouped similar metrics into the following nine metric types: count-based, diversity, activity-based, physiological, survival, reproductive success, education-based, regulation-based, and biomass (Appendix S2).

We quantified the number of studies per conservation intervention that met certain relevance and study design criteria, to give an estimate of the availability of relevant and robust evidence. To ensure that we did not artificially constrain the number of studies per intervention for different subsets of studies (e.g., taxonomic order or biome), we grouped certain interventions that were focused on single taxa or habitats but were fundamentally the same type of intervention (e.g., ‘create ponds for newts’ and ‘create ponds for toads’ would be grouped into ‘create ponds’; see Acknowledgements and Data for files describing these groupings). This resulted in a total of 71 and 226 interventions for amphibians and birds, respectively.

Using these interventions, we then undertook two analyses to quantify the availability of evidence under different scenarios: i) where we optimistically assume a given practitioner is interested in the most frequently studied local context; and ii) where we assume that a given practitioner is interested in local contexts in which a greater percentage of species are threatened (i.e., those classified as Vulnerable, Endangered or Critically Endangered status on the (IUCN, 2019) Red List).

The first analysis calculated the mean number of studies per intervention for both scenarios in terms of three separate relevance criteria: biome, taxonomic order and metric. For the first scenario we calculated the number of studies with the most frequently studied biome, order or metric relative to each intervention. For the second scenario (to reflect conservation needs), we calculated the number of studies with a randomly selected biome, taxonomic order or metric from a weighted list (averaged over 1,000 repeated runs). This weighted list was generated so that the probability of selection was determined by the percentage of species that are threatened (i.e., those classified as Vulnerable, Endangered or Critically Endangered status on the (IUCN, 2019) Red List) for each biome and taxonomic order, and the percentage usage of each metric within each intervention in the database. We intersected shapefiles from the (IUCN, 2019) Red List with shapefiles of the world’s terrestrial biomes (Dinerstein et al., 2017) to determine the proportion of threatened species in each biome. We assumed that interventions could be tested by studies in any biome and on any taxonomic order - this will likely mean that our estimates for the second scenario are underestimates of study availability, for example, as certain interventions are unlikely to be conducted in certain biomes. However, we grouped interventions so they were not defined as taxon or habitat-specific and used coarse criteria (biome and taxonomic order) to limit this underestimation.

For the second analysis, we used a stepwise process to calculate the number of studies that met one or more of the relevance criteria - only carrying forward studies if they met all previous criteria. For example, considering the first scenario (most frequently studied context), we counted the number of studies featuring the most frequently studied biome, then studies featuring the most frequently studied biome AND taxonomic order, and then studies featuring the most frequently studied biome AND taxonomic order AND metric. We also repeated this for all possible orderings of biome, taxonomic order and metric (Fig.3 and Figs.S1-S5), as well as for the second scenario (weighting towards biomes and taxonomic orders with greater percentages of threatened species). Taxonomic orders could only be selected if at least one species in that order was present in the previously selected biome - we determined which orders were present in each biome by intersecting shapefiles from the (IUCN, 2019) Red List with shapefiles of terrestrial biomes (Dinerstein et al., 2017). The same was true for biomes when taxonomic order was the first relevance criteria to be selected (i.e., only biomes where that taxonomic order is present could be selected). In the final step, we also calculated the number of studies that used different categories of study designs (any design; BA, CI, BACI or RCT; CI, BACI or RCT; BACI or RCT).

**Figure 2.**
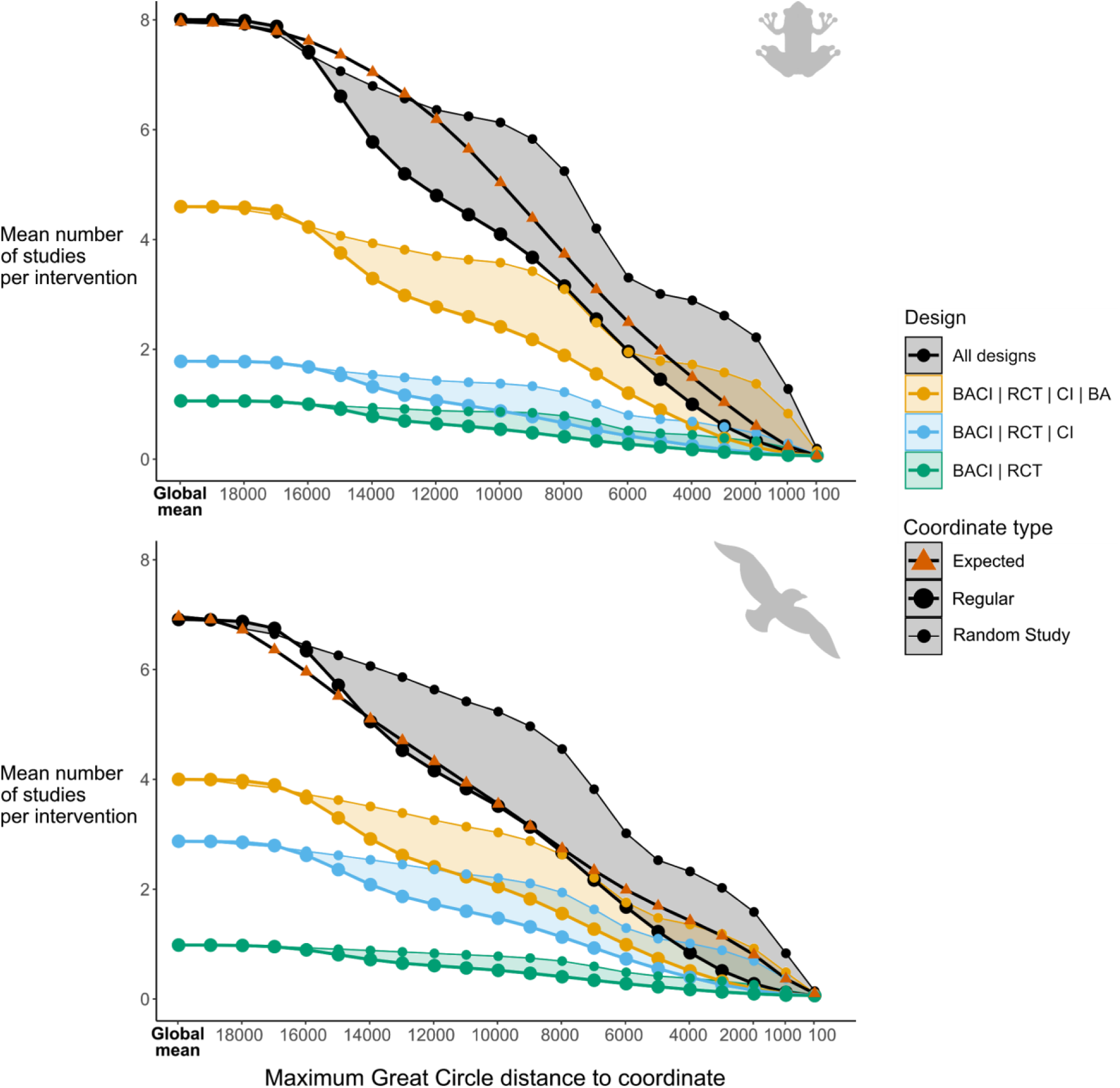
The mean number of amphibian and bird studies per intervention using different study designs found within a certain distance of different sets of coordinates. The maximum distance that a study can be is shown on the x axis, starting with the Global Mean (mean number of studies per intervention considering all studies in the database) and decreasing to a distance of 100 km. Regular coordinates (large circle, thick line) show the mean number of studies within a certain distance from a set of regularly distributed coordinates. Expected coordinates (orange triangle) mimic how the availability of studies would be expected to change if studies were regularly distributed (this is only shown for studies using any study design). Random Study coordinates (small circle, thin line) show the mean number of studies within a certain distance from a set of randomly selected coordinates where previous studies have been conducted.

**Figure 3.**
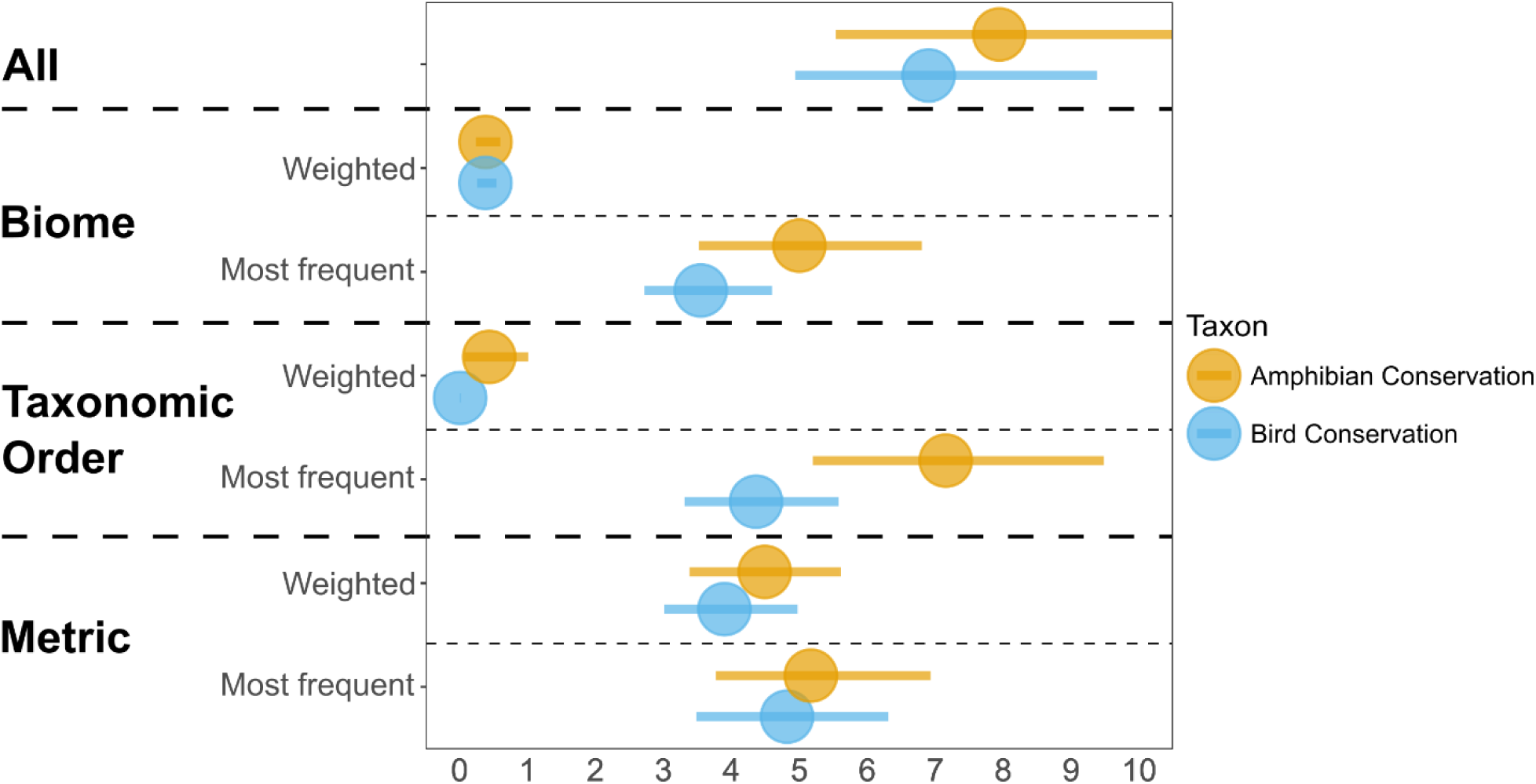
Mean number of studies per intervention when studies were counted based on whether they considered the most frequently studied biome, metric or order, and whether they considered a randomly selected biome, metric or taxonomic order from a weighted list. These weightings were based on the proportion of threatened species found in each biome or taxonomic order. ‘All’ indicates the mean number of studies per intervention when considering all studies.

## Results

We considered a total of 71 and 226 interventions for amphibians and birds (mean = 7.9 and 6.9 studies per intervention; Fig.2), respectively, that contained at least one study. Studies were not evenly distributed geographically; the mean number of amphibian and bird studies per intervention (black large circles in Fig.2) deviated, particularly for amphibians, from what we would have expected if the same number of studies were regularly distributed (orange triangles in Fig.2). On average, there was less than one study per intervention available within 2,000km from a given regular point. When restricting analyses to more robust designs, the availability of studies decreased substantially, with a higher proportion of amphibian studies using BA designs, compared to birds, but a smaller proportion using CI (see drop-offs from orange to blue, and blue to green lines, respectively; Fig.2).

When considering distance of studies to randomly selected study coordinates, the mean number of studies per intervention generally declined more gradually compared to a regular grid of coordinates (Fig.2), implying that studies are clustered in space. At distances below 5,000km these differences were particularly pronounced; for example, on average, 2.2 amphibian studies and 1.5 bird studies were within 2,000km of a random study coordinate, compared to only 0.3 amphibian studies and 0.2 bird studies within 2,000km of regularly spaced coordinate. This suggests that studies are slightly more clustered for amphibians than birds.

The mean number of studies per intervention was substantially greater for the most frequently studied biome (Amphibians: 5.0; Birds: 3.5), relative to each intervention, compared to biomes with higher percentages of species that are threatened (Amphibians: 0.4; Birds: 0.4; Fig.3). Similarly, the mean number of studies per intervention was substantially greater for the most frequently studied order in each intervention (Amphibians: 7.2; Birds: 4.4), compared to a taxonomic orders with higher percentages of species that are threatened (Amphibians: 0.4; Birds: 0.01; Fig.3). There was a smaller difference in the mean number of studies per intervention between studies that used the most frequently used metric (Amphibians: 5.2; Birds: 4.8), relative to each intervention, and studies that used a randomly selected metric from within each intervention (Amphibians: 4.5; Birds: 3.9; Fig.3). The mean numbers of biomes, taxonomic orders and metrics per intervention were 2.7, 2.6, and 3.1 for amphibians, respectively, and 2.4, 6.1, and 2.6 for birds, respectively.

The mean number of studies per intervention was also greater when we constrained by the most frequently studied biome, taxonomic order and metric in a stepwise process Fig.4A), compared to biomes and taxonomic orders with higher percentages of threatened species (Fig.4B). When we constrained by the most frequently studied biome, taxonomic order and metric, the greatest proportional decrease in the number of studies occurred once we further constrained by study design, by only counting studies using robust BACI or RCT designs (on average, ∼20% of amphibian studies and ∼17% of bird studies that had met all previous criteria; Fig.4A). When we constrained by biomes and taxonomic orders with higher percentages of threatened species, the greatest proportional decreases occurred when constraining by taxonomic order, most notably for birds, and by biome (Fig.4B).

**Figure 4.**
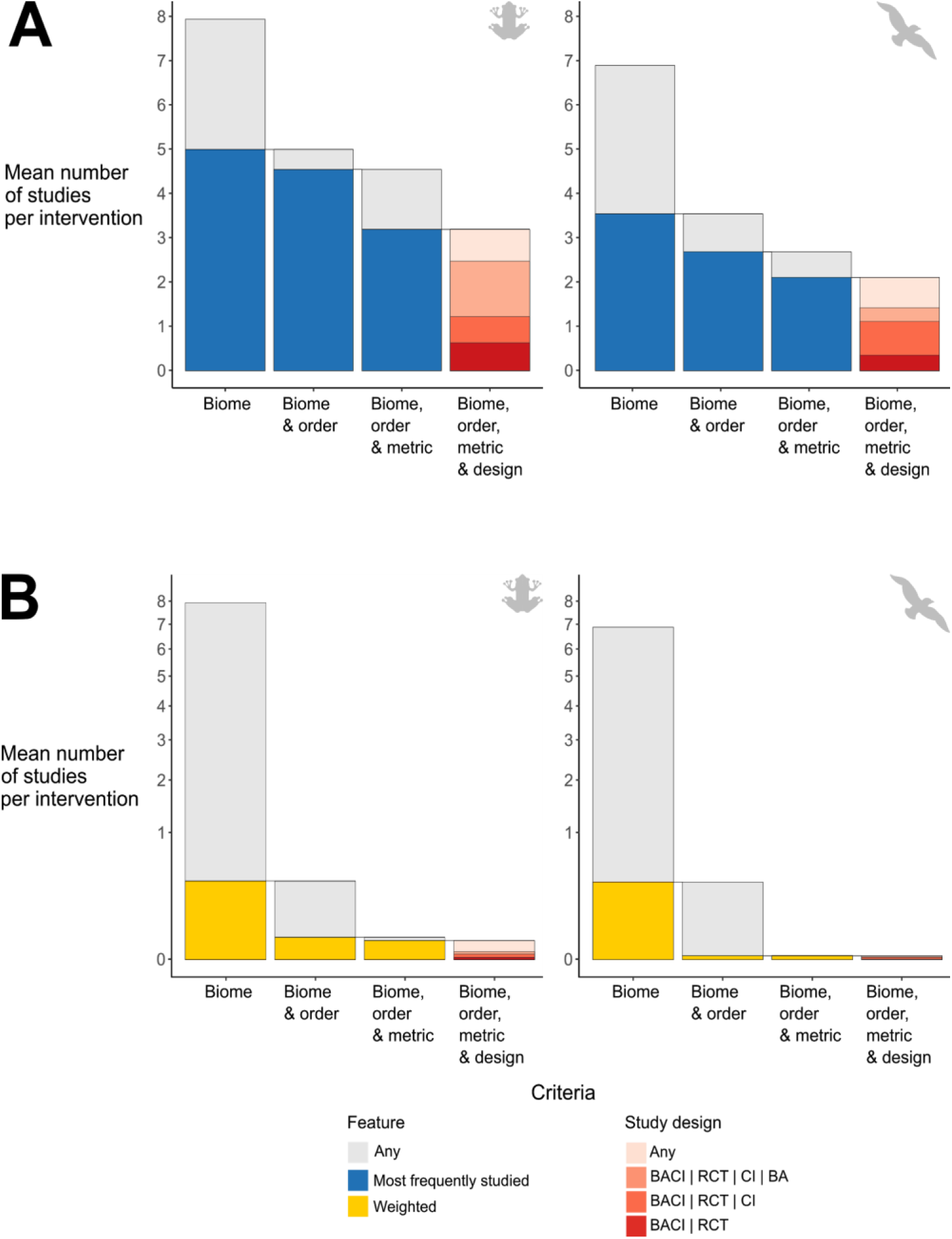
Mean numbers of amphibian and bird studies per intervention when only considering studies that meet certain relevance criteria. In panel A, studies with the most frequently studied biome, taxonomic order and metric relative to each intervention were counted - here we assume practitioners are interested in the most frequently studied local context. At each step (left to right) we add a further criterion, carrying forward relevant studies from the previous step - for example, only studies conducted in the most frequently studied biome were carried forward into the biome and order category. In panel B, studies with a selected biome, taxon and metric were counted (y axis has a square root transformation). Here we assume practitioners are more likely to be interested in: biomes that are inhabited by higher proportions of threatened species; taxonomic orders that have higher relative proportions of threatened species; and metrics that are most frequently used within each intervention. At the final step, studies are counted based on the study design they use (see Methods for details of study designs).

The sequence in which criteria were applied did not substantially affect the magnitude of the decrease in the number of studies - e.g., when biome was selected before or after taxonomic order and metric (Supporting Information Fig.S1-5). The overall decrease in studies from applying all relevance criteria (biome, taxonomic order and metric) was similarly severe regardless of the sequence in which the criteria were applied (Supporting Information Fig.S1-5). For all sequences, constraining the evidence to studies that used robust BACI or RCT designs reduced the mean number of studies to less than one study after constraining by the most frequently studied biome, taxonomic order and metric (Fig.4A; Supporting Information Fig.S1-5). Doing the same after instead constraining by the biomes and taxonomic orders with higher percentages of threatened species reduced the mean number of studies to fewer than 0.01 studies with BACI or RCT designs (Fig.4B; Supporting Information Fig.S1-5).

## Discussion

Our work demonstrates that not only is there a general paucity of studies testing conservation interventions, but that the distribution of these studies does not reflect conservation needs. Specifically, there is a lack of studies testing conservation interventions in biomes and for taxonomic orders containing high percentages of threatened amphibian and bird species. Given substantial declines of bird fauna (Rosenberg et al., 2019) and severe threats to amphibians (Grant, Muths, Schmidt, & Petrovan, 2019), a better understanding of the effectiveness of interventions targeting threatened species is urgently required. Furthermore, a given decision-maker is likely to struggle to find robust studies addressing their local context. Addressing this deficit will be challenging, but there are several possible ways to improve the evidence base for conservation.

A fundamental problem that needs to be overcome in the long-term is the lack of studies testing conservation interventions. Williams, Balmford, & Wilcove (*in review*) found that only 15% of studies from a representative sample of the conservation literature tested interventions. Evaluation of interventions should become mainstream, both as a topic of academic research and as an activity for on-the-ground conservationists (Baylis et al., 2016). The publication of these tests, whether the results are positive, negative, or neutral, is critical to building a strong evidence base for conservation (Catalano, Lyons-White, Mills, & Knight, 2019). Current efforts to facilitate this include the Applied Ecology Resources repository (British Ecological Society, 2020), ‘Evidence’ articles in the journal Conservation Science and Practice (Society for Conservation Biology, 2020), and the journal Conservation Evidence (Conservation Evidence, 2020b).

Simply publishing more tests of conservation interventions, even at an increasing rate, is however unlikely to solve the paucity of locally relevant studies. For example, even though adding 1,000 studies testing interventions on birds would increase the mean number of studies to approximately 11 studies across the current 226 interventions, these studies would still be spread thin across a myriad of local contexts where the need for conservation is often not the greatest (see also Wilson et al., 2016). Although Reboredo Segovia, Romano, & Armsworth (2020) suggest that the number of general conservation studies in tropical locations correlates with the number of threatened species, the results of this study and (Christie, Amano, Martin, Petrovan, et al., 2019) suggest this is not the case for conservation studies testing interventions. Therefore, we need concrete solutions enabling conservationists to generate and collate more experimental evidence on the effectiveness of conservation interventions in underrepresented locations and on underrepresented taxa (Christie, Amano, Martin, Petrovan, et al., 2019; Donaldson et al., 2016; Murray, Green, Williams, Burfield, & de Brooke, 2015). For example, funders, principal investigators and heads of conservation organizations need to enhance and prioritize funding to test interventions in underrepresented areas. Evidence synthesis also needs to incorporate more evidence from non-English language and grey literature publications to help address underrepresented local contexts (Amano, González-Varo, & Sutherland, 2016; Amano & Sutherland, 2013) - for example, publications from over 317 non-English language journals are starting to be added to the Conservation Evidence database through the Transcending Language Barriers to Environmental Sciences project (TRANSLATE, 2020). Making concerted efforts to acquire grey literature from organizations and groups outside academia will also be important.

The low proportion of studies using robust study designs, regardless of their relevance to a local context, is also challenging. That more robustly designed studies are concentrated in North America, Europe and Australia also compounds earlier taxonomic and biogeographical biases (Christie, Amano, Martin, Petrovan, et al., 2019). If few robustly designed studies are available for informing conservation, decision-makers may have to consider a wider range of studies that may be less robust or relevant, potentially reducing the effectiveness of decision-making and future practice (Slavin, 1995; Tugwell & Haynes, 2006; Whittaker, 2010). To increase the quality of studies available for decision-making, we must recognize that the quality of studies testing interventions may be limited in different ways. Studies evaluating mitigation efforts are often not constrained by cost, but rather by short timescales and their focus on meeting legislative requirements (for example, conserving legally protected species). Studies testing non-mitigation interventions will likely be more constrained by cost, as well as short timescales (e.g., PhD funding). Acknowledging how real-world constraints affect the choice of study design is essential to devising approaches to improving the evidence base for conservation. While better training of early career scientists, consultants and researchers in appropriate study designs for causal inference may help, ultimately more regulatory and funder-led measures (e.g., requiring grantees to demonstrate rigorous study design) will be required (De Palma et al., 2018; Grant et al., 2019).

Given the general lack of evidence across conservation, there is also a need to use a standardized set of metrics to evaluate conservation effectiveness (McQuatters-Gollop et al., 2019). Using a diversity of metrics may be necessary to assess multiple important aspects of an intervention’s effectiveness, but a lack of consistency in the metrics used to report results often makes the evidence base difficult to synthesize - especially if different metrics yield different results (Mace & Baillie, 2007). Prioritisation of the most relevant metrics of effectiveness for different interventions with input from decision-makers and practitioners is essential to facilitate inter-study comparisons (McQuatters-Gollop et al., 2019). Initiatives aiming to do this are underway in topics such as fishery habitats (Lederhouse & Link, 2016) and protected areas (Nolte & Agrawal, 2013; Pomeroy, Parks, & Watson, 2004), and are supported by the Essential Biodiversity Variables framework (Jetz et al., 2019). Funders could help strengthen these efforts by requiring grantees to follow such initiatives and use consistent metrics when evaluating interventions.

Increasing the size and quality of the evidence base for conservation decision-making will be a slow process, but conservation practitioners need to make decisions now. Until the evidence base improves, excluding studies from evidence syntheses because they do not meet certain quality or relevance criteria could lead to little or no evidence being used to inform conservation efforts (Davies & Gray, 2015; Gurevitch & Hedges, 1999; Lortie, Stewart, Rothstein, & Lau, 2015). Moreover, studies that do not meet these criteria may still provide useful evidence, particularly in the absence of more relevant and robust studies (Burivalova et al., 2019; Cook, Mascia, Schwartz, Possingham, & Fuller, 2013; Gough & White, 2018).

Therefore, we need novel approaches to rigorously synthesizing studies that vary considerably in their relevance and robustness to maximize the use of the current imperfect evidence base. We believe that weighting approaches in both quantitative meta-analyses and more qualitative evidence synthesis would help maximize the number of studies available, while giving greater influence to studies with desirable characteristics. This could involve giving greater influence to more robustly designed studies (e.g., using accuracy weights from Christie, Amano, Martin, Shackelford, et al. 2019 and evidence hierarchies from Mupepele, Walsh, Sutherland, & Dormann 2016), and giving more weight to more relevant studies (e.g., weighting by the relevance of studies to a decision-maker’s local context, as proposed in healthcare by Kneale, Thomas, O’Mara-Eves, & Wiggins 2019). To generate objective weights of study relevance that reflect the likely generalizability of study results, we need studies which help us to understand how generalizability varies between interventions for different ecological (e.g., artificial nest boxes; Finch et al. 2019), socioeconomic, and political contexts. Understanding why some interventions work in certain contexts and not others is fundamentally important for evidence-based decision-makers (Grant et al., 2019).

Overall, we have shown that the evidence base for conservation does not reflect the needs of conservation. When this is combined with the general paucity of robust studies testing conservation interventions, we conclude that there is a serious lack of locally relevant and robust studies to inform decision-making in conservation. We hope that the conservation community can work together to improve the state of the conservation evidence base. Doing so will require much greater collaboration between research and practice. Testing interventions needs to become more routine, use a more standardized suite of metrics and robust study designs, and, most importantly, focus on the locations and taxa where evidence is most needed to inform conservation action. In the meantime, we need to explore ways to better analyze the current patchy evidence base of conservation and ensure that we can support the shift towards more evidence-based policy and practice.

## Acknowledgements and Data

We would like to thank Anne Mupepele for their useful comments on the manuscript and all past and present members of the Conservation Evidence project. All data analyzed in this study and code to repeat analyses are available from https://doi.org/10.5281/zenodo.3634780.

