## Supporting Information for "Poor availability of context-specific evidence hampers decision-making in conservation"

Affiliations:
[1]Conservation Science Group, Department of Zoology, University of Cambridge, The David Attenborough Building, Downing Street, Cambridge CB3 3QZ, UK.
[2] BioRISC, St. Catharine's College, Cambridge CB2 1RL, UK.
[3]School of Biological Sciences, University of Queensland, Brisbane, 4072 Queensland, Australia.
[4] Department of Animal and Plant Sciences, University of Sheffield, Sheffield, S10 2TN United Kingdom.
[5]Centre for Ecology and Conservation, College of Life and Environmental Sciences, University of Exeter, Penryn, UK
[6]Sustainability Research Institute, School of Earth and Environment, University of Leeds. LS2 9JT United Kingdom

### Figures S1-S5

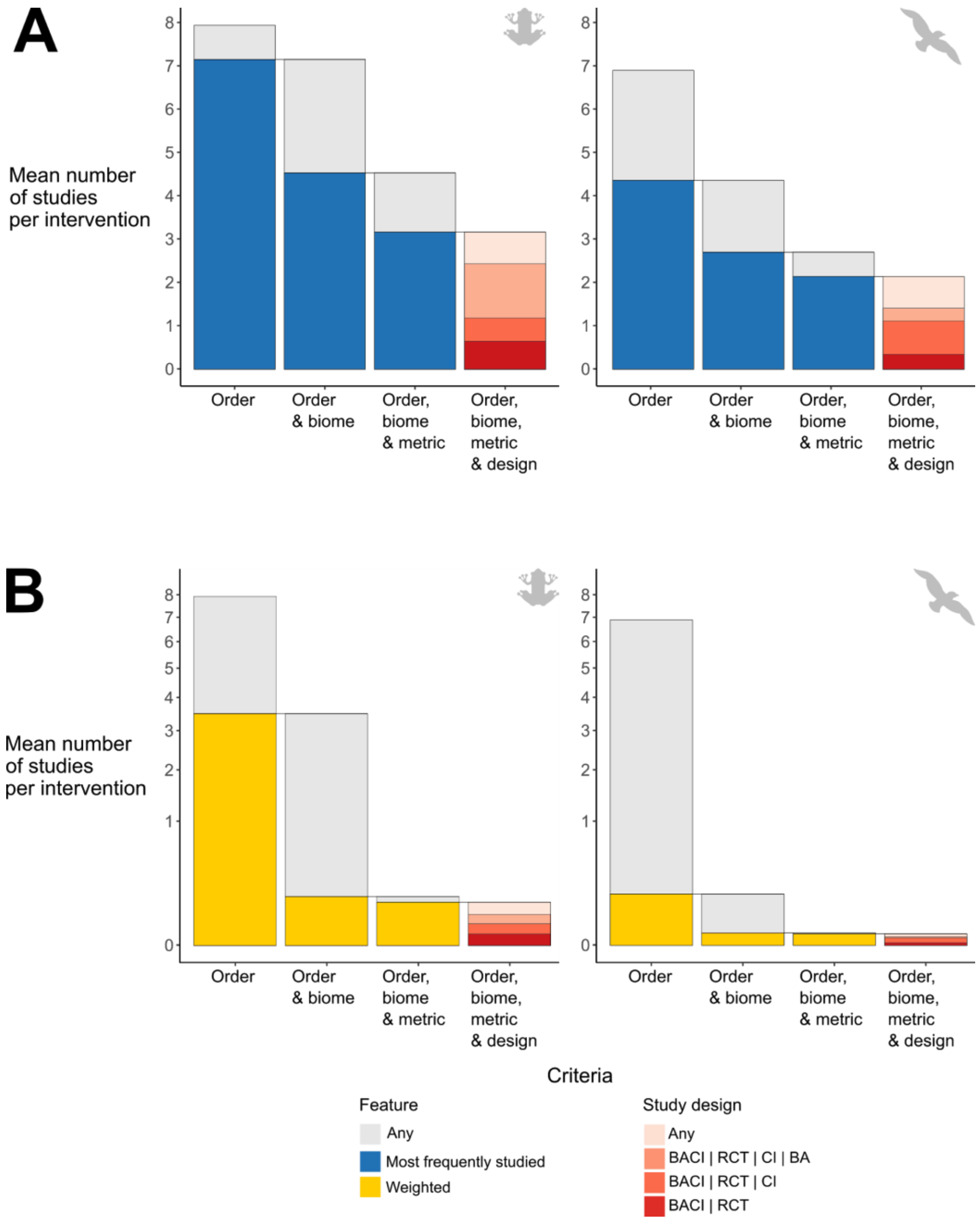


Figure S1 - Mean numbers of amphibian and bird studies per intervention when only considering studies that meet certain relevance criteria. In panel A, studies with the most frequently studied taxonomic order, biome and metric relative to each intervention were counted - here we assume practitioners are interested in the most frequently studied local context. At each step (left to right) we add a further criterion, carrying forward relevant studies from the previous step - for example, only studies with the most frequently studied order were carried forward into the order and biome category. In panel B, studies with a selected taxonomic order, biome and metric were counted (y axis has a square root transformation). Here we assume practitioners are more likely to be interested in: biomes that are inhabited by higher proportions of threatened species; taxonomic orders that have higher relative proportions of threatened species; and metrics that are most frequently used. At the final step, studies are counted based on the study design they use (see Methods for details of study designs).

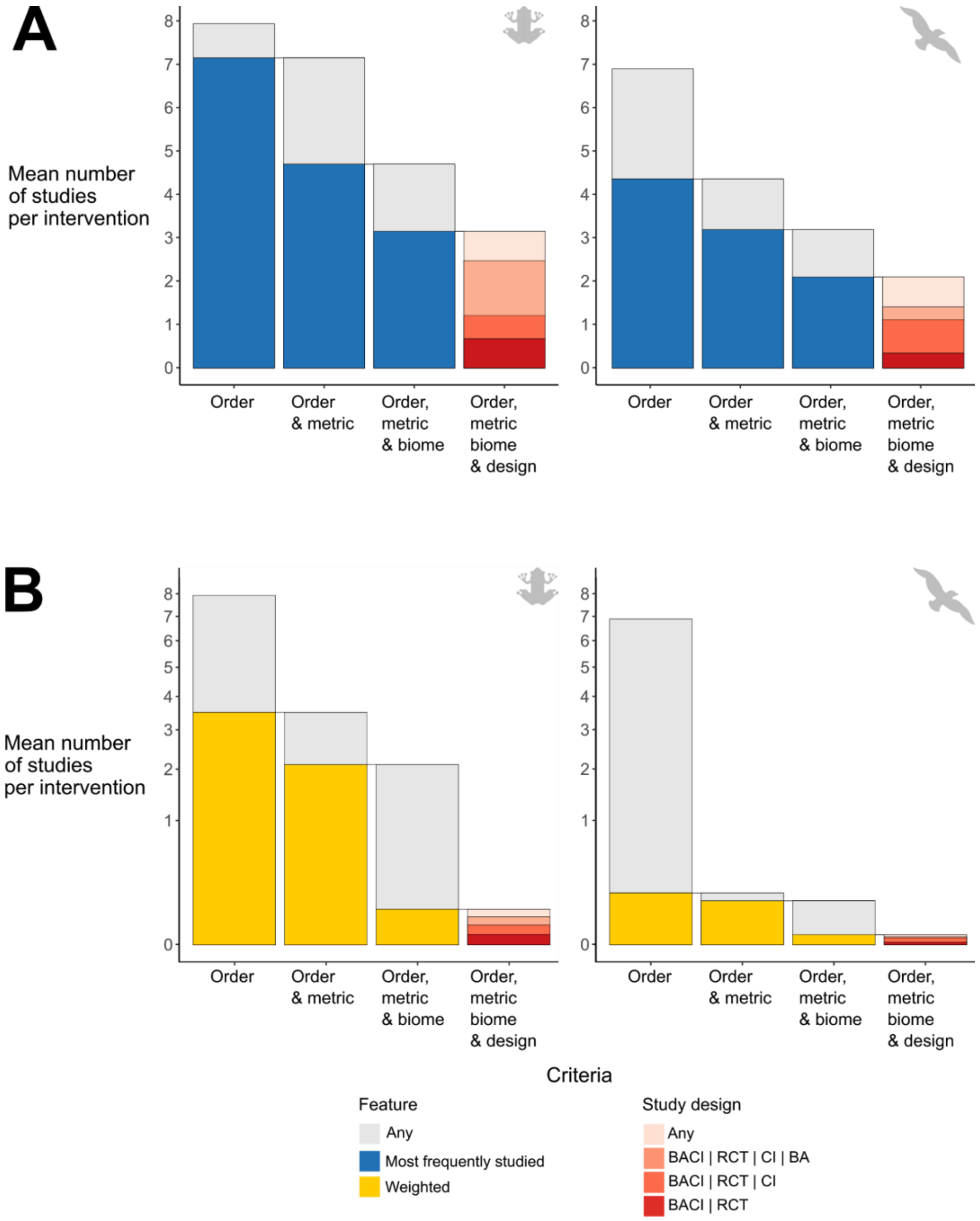


Figure S2 - Mean numbers of amphibian and bird studies per intervention when only considering studies that meet certain relevance criteria. In panel A, studies with the most frequently studied taxonomic order, metric and biome relative to each intervention were counted - here we assume practitioners are interested in the most frequently studied local context. At each step (left to right) we add a further criterion, carrying forward relevant studies from the previous step - for example, only studies with the most frequently studied order were carried forward into the order and metric category. In panel B, studies with a selected taxonomic order, metric and biome were counted (y axis has a square root transformation). Here we assume practitioners are more likely to be interested in: biomes that are inhabited by higher proportions of threatened species; taxonomic orders that have higher relative proportions of threatened species; and metrics that are most frequently used. At the final step, studies are counted based on the study design they use (see Methods for details of study designs).

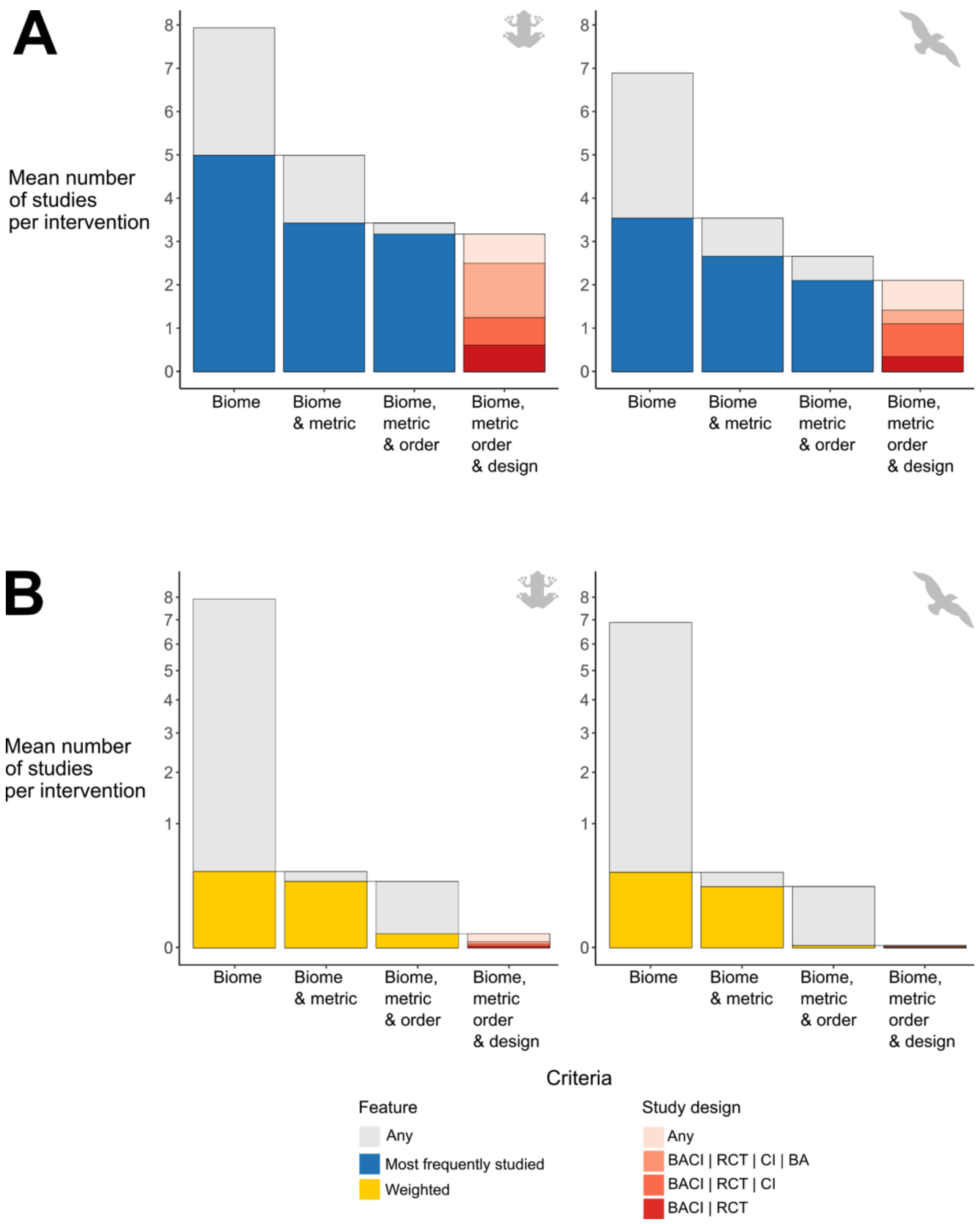


Figure S3 - Mean numbers of amphibian and bird studies per intervention when only considering studies that meet certain relevance criteria. In panel A, studies with the most frequently studied biome, metric and taxonomic order relative to each intervention were counted - here we assume practitioners are interested in the most frequently studied local context. At each step (left to right) we add a further criterion, carrying forward relevant studies from the previous step - for example, only studies with the most frequently studied biome were carried forward into the biome and metric category. In panel B, studies with a selected biome, metric and taxonomic order were counted (y axis has a square root transformation). Here we assume practitioners are more likely to be interested in: biomes that are inhabited by higher proportions of threatened species; taxonomic orders that have higher relative proportions of threatened species; and metrics that are most frequently used. At the final step, studies are counted based on the study design they use (see Methods for details of study designs).

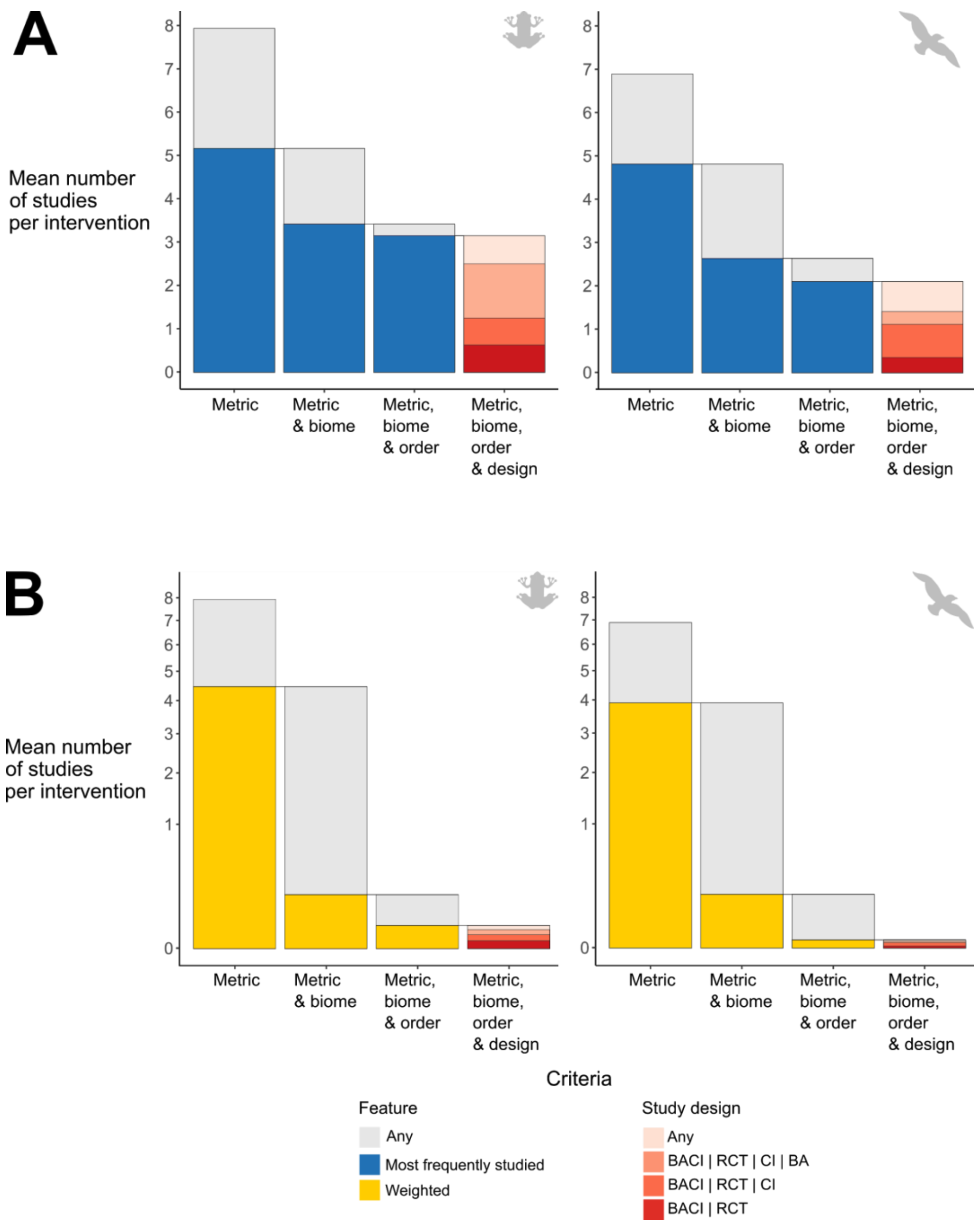


Figure S4 - Mean numbers of amphibian and bird studies per intervention when only considering studies that meet certain relevance criteria. In panel A, studies with the most frequently studied metric, biome and taxonomic order relative to each intervention were counted - here we assume practitioners are interested in the most frequently studied local context. At each step (left to right) we add a further criterion, carrying forward relevant studies from the previous step - for example, only studies with the most frequently studied metric were carried forward into the metric and biome category. In panel B, studies with a selected metric, biome and taxonomic order were counted (y axis has a square root transformation). Here we assume practitioners are more likely to be interested in: biomes that are inhabited by higher proportions of threatened species; taxonomic orders that have higher relative proportions of threatened species; and metrics that are most frequently used. At the final step, studies are counted based on the study design they use (see Methods for details of study designs).

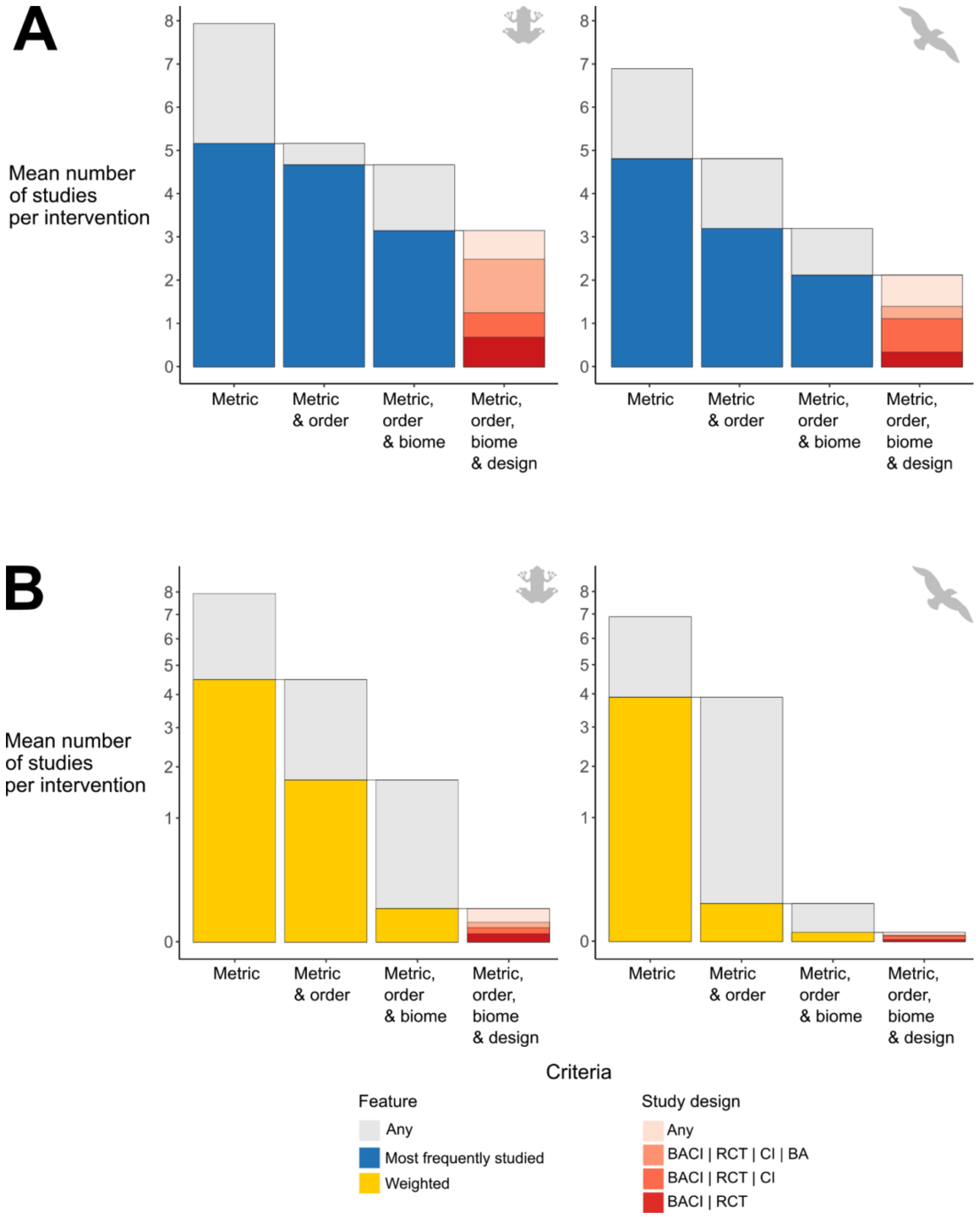


Figure S5 - Mean numbers of amphibian and bird studies per intervention when only considering studies that meet certain relevance criteria. In panel A, studies with the most frequently studied metric, taxonomic order and biome relative to each intervention were counted - here we assume practitioners are interested in the most frequently studied local context. At each step (left to right) we add a further criterion, carrying forward relevant studies from the previous step - for example, only studies with the most frequently studied metric were carried forward into the metric and order category. In panel B, studies with a selected metric, taxonomic order and biome were counted (y axis has a square root transformation). Here we assume practitioners are more likely to be interested in: biomes that are inhabited by higher proportions of threatened species; taxonomic orders that have higher relative proportions of threatened species; and metrics that are most frequently used. At the final step, studies are counted based on the study design they use (see Methods for details of study designs).

### Appendix S1

To generate regularly spaced terrestrial coordinates for birds and regularly spaced coordinates within the combined extent of all amphibian species ranges, we used R statistical software version 3.5.1 (R Core Team, 2019) and the packages sp (R. S. Bivand, Pebesma, & Gomez-Rubio, 2013; Pebesma & Bivand, 2005), rgdal (R. Bivand, Keitt, & Rowlingson, 2019) and rgeos (R. Bivand & Rundel, 2019). We first generated a regularly spaced grid of coordinates, checked which coordinates fell within the appropriate shapefiles (from "OpenStreetMap" 2019 for birds and the IUCN 2019 for amphibians), and adjusted until we produced the desired number of regularly spaced coordinates - R code to perform all analyses is available at https://doi.org/10.5281/zenodo.3634780.. Maps of regularly spaced coordinates for amphibians and birds are presented below. To calculate the Great Circle Distance from each study to each coordinate, we used the geosphere package (Hijmans, 2017) with R statistical software version 3.5.1 (R Core Team, 2019).

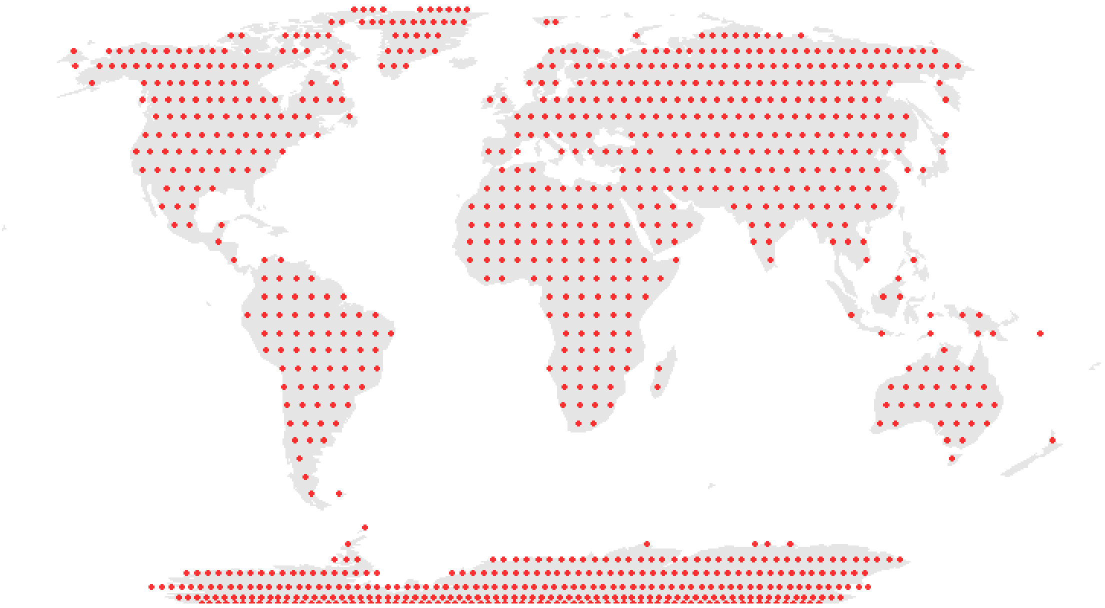

Fig.S6 - Regularly spaced coordinates for birds over terrestrial landmasses (OpenStreetMap 2019).

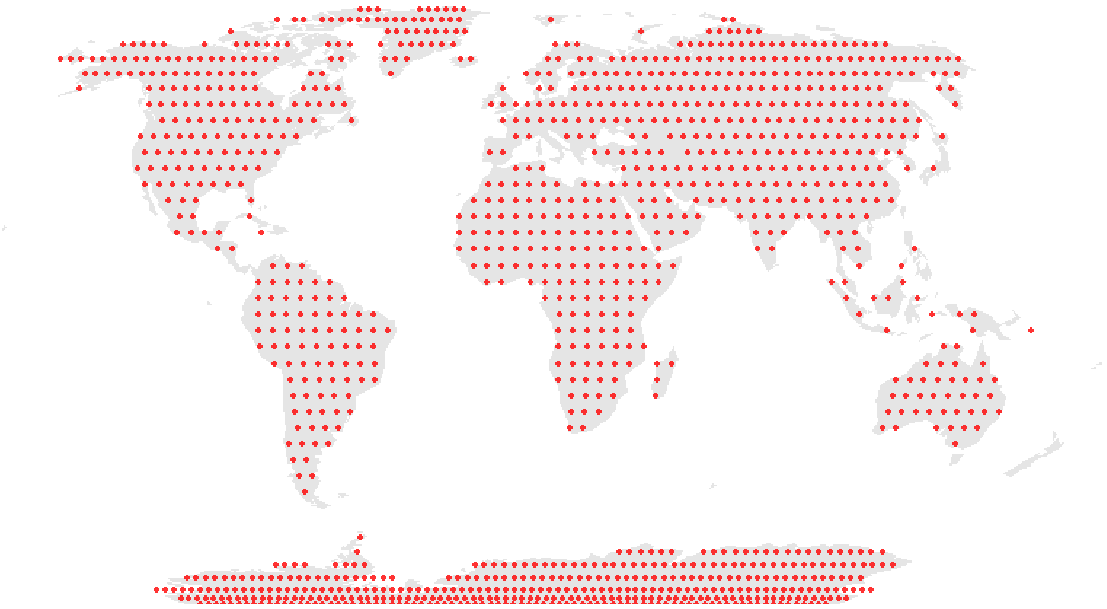

Fig.S7 - Expected coordinates for bird studies if studies were regularly distributed over terrestrial landmasses (OpenStreetMap 2019).

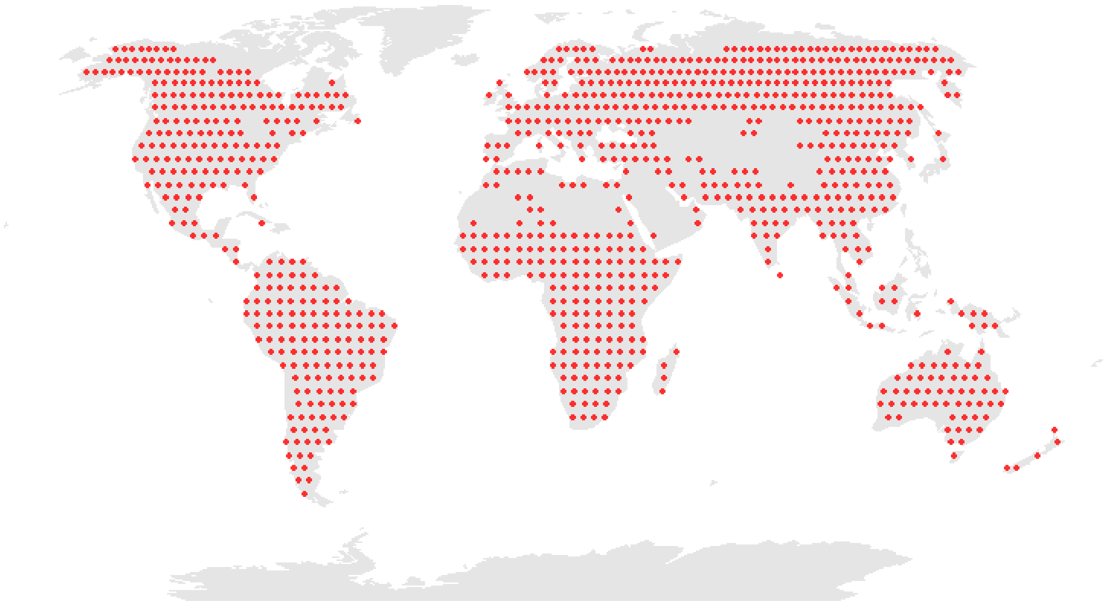

Fig.S8 - Regularly spaced coordinates for amphibians over the combined extent of all amphibian species ranges (IUCN 2019).

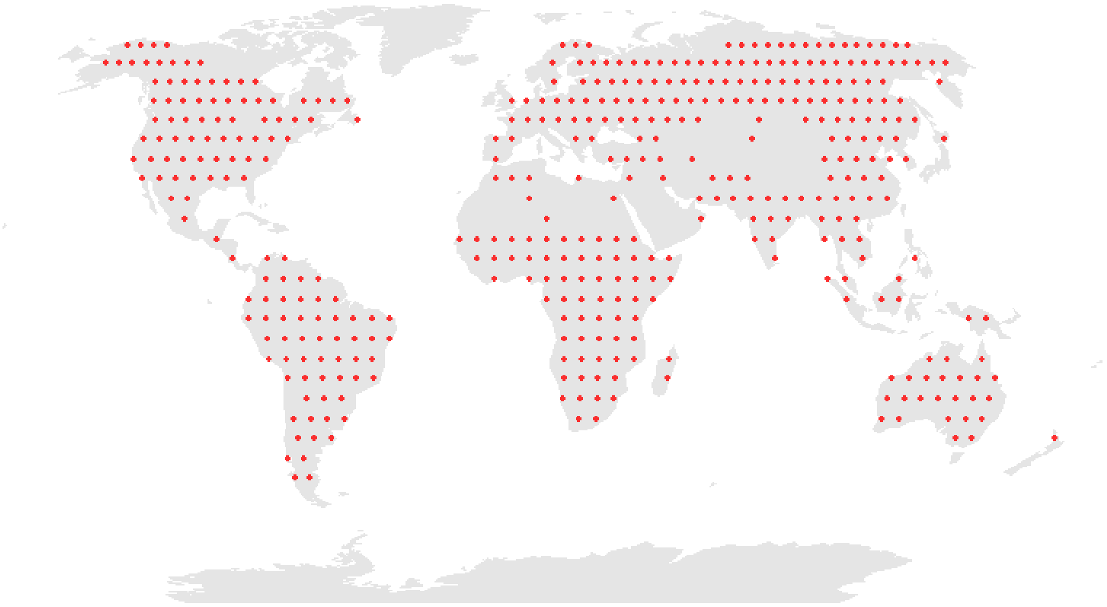

Fig.S9 - Expected coordinates for amphibian studies if studies were regularly distributed over the combined extent of all amphibian species ranges (IUCN 2019).

### Appendix S2

To obtain the metadata we needed to assess the availability of relevant studies, we used two previously described methods from Christie, Amano, Martin, Petrovan, et al. (2019): i) assigning each study to a biome using coordinates from the Conservation Evidence database, the sp package in R (Bivand, Pebesma, & Gomez-Rubio, 2013; Pebesma & Bivand, 2005) and a shapefile obtained from Dinerstein et al. (2017) - see https://doi.org/10.5281/zenodo.3634780 for R code; and ii) web-scraping of the Conservation Evidence website to obtain the metrics and study design used by each study (see Christie, Amano, Martin, Petrovan, et al. 2019 for R code for extracting study design and https://doi.org/10.5281/zenodo.3634780 for R code to extract metrics). In Christie, Amano, Martin, Petrovan, et al. (2019), we only considered four broad metric types (abundance/density/cover, reproductive success, diversity and survival/mortality), but here we expanded this, extracting 14 different metrics from study summaries on the Conservation Evidence website, which we grouped into the following nine groups: count-based (abundance, density and cover), diversity, activity-based (activity, frequency of usage and occupancy), physiological, survival (survival and mortality), reproductive success, education-based, regulation-based, and biomass. Details of keywords used to extract metadata (adapted from methods in Christie, Amano, Martin, Petrovan, et al. 2019) are found in the R code for extracting metrics available at https://doi.org/10.5281/zenodo.3634780. These reflected similar broad types of metrics that may be used to assess the effectiveness of an intervention to conserve birds or amphibians. The accuracy of metric type extraction from the Conservation Evidence website was 86% from a random 5% of amphibian studies (18 out of 21) and 90% for a random 5% of bird studies (56 out of 62) in the Conservation Evidence database. Accuracy was defined as there being no false positives or false negatives for that study in any intervention that it provided evidence for (as a single study can be found in multiple interventions).
